# OmniShim - a vendor-independent B0 Shimming software toolbox

**DOI:** 10.1101/2025.08.26.672438

**Authors:** Mahrshi Jani, Shengyue Su, Yeison Rodriguez, Andrew Wright, Kimberly Chan, Manoj Kumar Sarma, Sheeba Anteraper, Anke Henning

**Affiliations:** Advance imaging research Center. University of Texas Southwestern Medical center (UTSW), Dallas, TX, USA

**Keywords:** B_0_ shimming, ultrahigh field strengths, constrained fitting, single-voxel, image-based B_0_ shimming, magnetic resonance spectroscopic imaging, region of less interest

## Abstract

**Purpose:** To introduce the OmniShim Toolbox, a software package designed to calibrate and perform B_0_ shimming in user-defined regions of interest across systems from different vendors, supporting various shim orders.

**Methods:** In this study, we systematically compared the performance of vendor-implemented B_0_ shim routines with our custom developed OmniShim toolbox, designed to improve static magnetic field homogeneity. Single-voxel magnetic resonance spectroscopy (MRS) data were acquired from the prefrontal cortex and occipital lobe, while magnetic resonance spectroscopic imaging (MRSI) data were collected from regions above and below the corpus callosum in healthy volunteers. Measurements were conducted on both 3T and 7T MR systems to evaluate the robustness and scalability of each B0 shimming strategy across different field strengths. Additionally, functional MRI (fMRI) data were acquired at 7T to assess the impact of improved shimming on EPI data quality and BOLD contrast.

**Results:** The OmniShim Toolbox demonstrated superior B_0_ homogeneity across all applications and field strength, leading to reduced signal dropout in fMRI and MRSI data, significantly improved spectral linewidths in SV MRS and MRSI data as well improved detection of neural networks by resting state fMRI.

**Conclusion:** The proposed OmniShim Toolbox offers a robust and flexible approach to control B_0_ inhomogeneity, resulting in substantial improvements in image quality and spectral resolution. These enhancements benefit a broad range of MR applications.

## INTRODUCTION

Magnetic resonance imaging (MRI) and magnetic resonance spectroscopy (MRS) techniques have a wide range of application in clinics and research. At high field strength we can detect more metabolites with proton and non-proton MRS and magnetic resonance spectroscopic imaging (MRSI) and achieve higher signal-to-noise ratios, higher spatial and spectral resolutions^1^. In addition, functional MRI specifically benefits from high magnetic field scanners by enhanced contrast-to-noise ratio and higher spatio-temporal resolution ^2^. However, to get full benefits of high field human MRI systems, B_0_ homogeneity is of critical importance for MRS and fMRI. B_0_ field inhomogeneities are induced by the subject under study, as air - tissue interfaces and interfaces between different tissue types distort the magnetic field according to their magnetic susceptibility, shape, and orientation relative to static magnetic field the MRI magnet produces. B_0_ field inhomogeneity introduces several artifacts in MRI data such as geometric distortion, loss of signal-to-noise ratio, signal dropout, and loss of spectral resolution in spectroscopy applications. Additionally, the amplitude of B_0_ inhomogeneities increases in a linear fashion with the strength of the magnetic field ^3–5^.

Active B_0_ shimming is usually performed by measuring the magnetic field in each individual subject and calculating the required shim currents applied to a set of B_0_ shim coils to counteract inhomogeneous B_0_ fields ^6–9^. Conventionally, these shim coils are designed to generate spherical harmonic field distributions ^7, 8^, as every shim field can be represented by a spherical harmonic basis function:

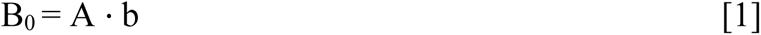

where, B_0_ is the given field distribution, A represents the distribution of harmonic function of fields, b is the amplitude of field strengths. Ideally, we need a robust, numerically stable, automated B_0_ shimming algorithm considering non-rectangular anatomically shaped regions of interest (ROI), while the B_0_ shim problem becomes ill conditioned due to shim current constraints especially in small ROIs and non-orthogonality in real B_0_ shim fields ^10,11^.

B_0_ shimming approaches can be classified into two categories: projection and volumetric mapping methods. Projection mapping methods like FASTERMAP, FASTESTMAP, and FASTMAP algorithms use generally rectangular shim volumes, and several linear projections passing through it, while neglecting all information outside of the respective beams ^12–16^. While there is consensus that projection based B_0_ shimming is a suitable method for single voxel MRS, ^17^the sparsity of the beams make them less suitable in case of either disjoint or large regions with complex geometries of interest such as whole-brain MRSI or fMRI or B_0_ shimming in the body. Alternatively, volumetric image-based B_0_ mapping provides a full FOV 3D field map but require longer acquisition time. Volumetric B_0_ shimming methods have been demonstrated to effectively improve field homogeneity within small regions of interest as well as in other body applications such as spinal cords etc. (ROIs; ^15, 18, 19^). Since anatomical structures rarely conform to simple rectangular geometries, the use of nonrectangular shim volumes has proven particularly advantageous ^15, 18–20^. However, excessively large B_0_ inhomogeneities outside the shimmed volume can induce artifacts inside the ROI ^17,18^. This has been addressed by defining two circumferent ROIs: an inner one corresponding to the target ROI and an outer one weighted lower in the cost function of the shim optimization problem to control artefacts (ROLI (region of less interest))^21^.

In high-field MR scanners, the number of B_0_ shim coils is often increased compared to lower field strengths, for better compensation of higher-order field components ^18–20, 22–24^. However, it has been shown that 2^nd^ and 3^rd^ SH shim fields delivered by in-bore vendor provided B_0_ shim systems deviate substantially from ideal SH B_0_ shim fields ^10, 11^. When the target shim volume is positioned away from the magnet’s isocenter, even ideal higher-order spherical harmonic terms may lose their inherent orthogonality, since spherical harmonics remain strictly orthonormal only within a spherical region centered at the isocenter. In addition, practical shim current limitations in combination with limited shim coil sensitivities lead to a restriction of the available maximum shim fields. These phenomena introduce additional complexity into determining localized shimming solutions since the shim current optimization problem becomes ill-conditioned and constraint. Hence, the non-orthogonality of real higher-order shim fields and the maximum shim field strength must be carefully addressed when computing the optimal currents for each shim coil. Moreover, choosing an appropriate B_0_ shimming algorithm with robust convergence becomes critical, as it needs to account for real shim fields and remain within hardware limits to effectively optimize the shim parameters for each unique imaging scenario ^10, 11, 25^ .

In this work, we present *OmniShim*, an image-based, vendor-independent B_0_ shim tool designed to improve the correction of magnetic field inhomogeneities. It considers 3D B_0_ maps as input providing full spatial coverage of the region of interest (ROI). Shim field impurities are considered by measuring real shim fields and deriving a cross-term calibration matrix that will be used during the B_0_ shim optimization ^10, 11^. An optimization algorithm with robust convergence in presence of the actual shim current constraints is implemented ^21, 25^. Flexible ROI selection options include read-in of rectangular single voxel MRS ROIs as planned on vendor’s console, automatic delineation of a whole-brain ROI by brain extraction, hand-drawn ROIs as well as the definition of two shim ROIs to consider the main target region and a region around it that may lead to artefacts and is considered in the shim algorithm with a lower weight in the cost function. We have evaluated our OmniShim tool at 3T MRI scanners from two different vendors and one 7T MRI scanner for B_0_ shimming problems in the head. Additionally, we compared the performance of vendor-supplied B_0_ shim algorithms with our OmniShim tool in both MRI and MRS applications across various brain regions at 3T and 7T

## SOFTWARE

### Software tool

The OmniShim software tool was developed in MATLAB (Natick, MA, USA). It uses dicom files of anatomical images and B_0_ maps as input and a text file with shim values as output. The output text file produced by the shimtool is directly ingestible by Philips MRI systems, whereas interfacing the shimtool with Siemens scanners requires a concise Bash script to facilitate the necessary communication. Details of the graphical user interface (GUI) of OmniShim is shown in supplemental figures 2-6. It provides visualization of the ROI position and B_0_ fields in the ROI before and after B_0_ shimming. OmniShim is designed to optimize B_0_ shimming for magnetic resonance imaging (MRI) by leveraging a combination of real shim field calibration ^10,11^, a robust shim algorithm in presence of shim field constraints, flexible ROI definition options and the FSL (FMRIB Software Library) ^26–28^ toolset. More specifically, OmniShim utilizes the FSL ‘prelude’ package for phase unwrapping of the B_0_ input maps and the ‘bet’ (brain extraction tool) for precise brain mask extraction, thereby enabling accurate and fast brain ROI definition for effective B_0_ shimming. The OmniShim tool is modular and can be easily integrated with multiple different vendor-integrated B_0_ shim systems and additional external B_0_ shim hardware.

**Figure 1** shows the OmniShim tool workflow, which begins by acquiring a three-dimensional gradient-echo dataset to generate a B_0_ field map, which serves as the fundamental input for shim current optimization. The B_0_ maps can be freely configured at the respective scanner consoles with respect to field-of-view, spatial resolution and delta TE to optimally consider different brain and body parts and achieve in-phase acquisition of water and fat ^30^ By default, the entire (unmasked) B_0_ field map is used, but an optional brain-extracted version—obtained via FSL’s BET algorithm is also available if one wishes to constrain the shim solution more specifically to brain tissue. From there, the user can define multiple regions of interest (ROIs) for B_0_ shimming, either via standard geometric shapes (using scanner voxel or field of view dimensions, off center and angulation parameters) or through manually specified slices and masks. This flexibility enables targeting of brain or body regions, while also allowing for adjustments to mitigate inhomogeneity in areas outside the primary ROI by selection of a secondary shim ROI. The tool also allows for slice-by-slice shimming. Full detail about ROI definition is given in the ROI definition paragraph below. By including these features in a streamlined interface, the OmniShim tool provides a robust, customizable method for refining B_0_ uniformity and addressing site-specific needs in a range of neuroimaging as well as clinical brain and body MRI applications.

**Figure1.**
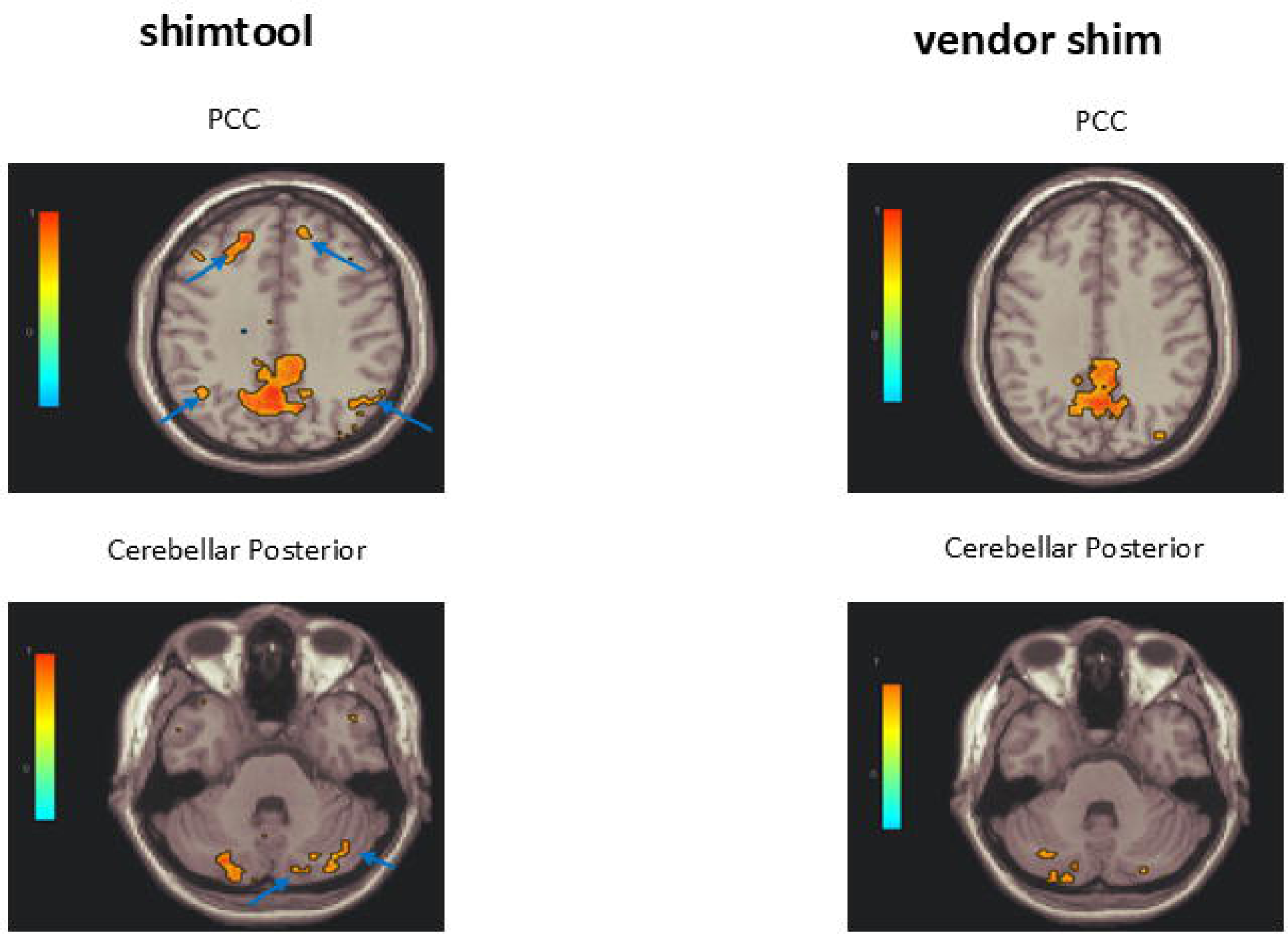
A) and (B) show the single voxel shimming comparison of vendor shimming with locally developed shimtool in the prefrontal cortex and visual cortex region of the brain. figures on the right show the comparison of calculated fullwidth half max (FWHM) of water linewidth. We have performed this experiment on our 3T system. Here we have compared shim performance with 2^nd^ order default shimming with our locally developed shimtool.

### Shim field calibration

The core of the OmniShim tool’s functionality lies in its ability to perform a comprehensive calibration of the real shim fields of the B_0_ shim coil array at hand. This process entails the establishment of a calibration matrix that fully characterizes the polarity and interactions among the shim fields of all B_0_ shim coils, thus ensuring accurate estimation and correction of the magnetic field distortions across the brain based on real shim fields ^10,11^. The calibration matrix is derived from high-resolution 3D B_0_ maps of phantoms obtained from each specific MRI scanner, which serve as the initial input for the tool. These maps provide detailed information about the shim field distribution of each shim coil across the imaging volume, serving as a critical basis for subject specific shim optimization. Notably, this approach is highly adaptable, as it requires only a one-time system-specific calibration at each MRI system. Once this calibration is completed, the method can be applied across different volunteers or patients, body parts and shim ROIs without the need for recalibration. Due to the scanner specific nature of the one-time calibration OmniShim is applicable to MRI systems from various vendors providing significant utility in multi-vendor environments. This ability to generalize across different MRI systems and field strength is a key strength, as it ensures that the B_0_ shimming process remains robust and reliable regardless of the scanner platform in use.

### Shim algorithm

Before calculating optimal shim solutions for different applications, phase unwrapping is applied to the input B_0_ maps of each masked region of interest using custom made phase unwarping solution so that phase wraps in phase of B_0_ maps do not impact shim optimization. The subject and ROI specific B_0_ shimming optimization is then performed through a pseudo-inverse based approach ^30–32^ to solve the B_0_ shim problem (equation 1). Singular value decomposition (SVD) is employed within this iterative procedure to adjust the shim currents in each shim coil in a way that the resulting magnetic field is as homogenous as possible across the ROI. In order, to respect the shim field constraints of the respective MRI system introduced by constraints in the shim field sensitivities and the maximum shim currents for each shim coil, the optimization process involves truncating singular values at each iteration^21^, progressively reinverting the matrix until the shim currents fall within the allowable limits set by the MRI system’s capabilities. This truncation approach ensures that the shim currents are both physically feasible and effective at minimizing B_0_ distortions ^22^. The pseudo-inverse approach is also robust against noise or localized artefacts in the B_0_ maps preventing the need of quality assurance of B_0_ maps or even manual exclusion of local artefacts as suggested earlier ^20^. The modular software approach allows for integration of alternative B_0_ shim optimization routines.

### ROI selection

<u>The OmniShim</u> tool offers flexible shim ROI selection options with freely chosen off-centers and angulations for MRI and MRS applications. Shim ROIs are generally not restricted to rectangular volumes, which is typically the case on all vendors scanner consoles. The user can define the shim ROI in three different ways: (i) rectangular ROIs defined by voxel or field-of-view (FOV) dimensions, angulations and off-centers of single voxel MRS ROIs or MRI FOVs, (ii) freely hand drawn 3D shim ROIs to allow for anatomically driven shim ROIs or (iii) automatic brain extraction for full-brain B_0_ shimming without needing to manually specify shim ROIs. This flexibility enhances the user experience and provides greater control over the B_0_ shimming process, allowing it to be tailored to specific regions of interest within the brain and body.

There are three advanced options related to defining flexible shim ROIs.

#### (1) Optional Brain Extraction

When enabled, brain extraction restricts the shim ROI overlayed on the input B_0_ map to intracranial tissue, excluding extracranial regions like the skull and scalp. This may improve shimming accuracy for correcting in-brain inhomogeneities, particularly in functional or structural MRI sensitive to off-resonance effects. This approach also provides a user independent and fast shim ROI definition for clinical brain imaging and neuroimaging research.

#### (2) Multiple Region-of-Interest

This advanced feature allows for considering a second ROI outside the primary shim ROI - called region of less interest (ROLI) ^21^ – which is considered but weighted lower than the primary ROI during the shim current computation. This approach is beneficial if certain peripheral areas are known to contribute to overall B_0_ field inhomogeneities like in cardiac imaging or spectroscopy or cause artefacts if not considered during the shim process like the skull lipid in brain MRSI. Supplemental Figure 5 illustrates the definition of ROI and ROLI for brain shimming; the advantage of this option was demonstrated previously ^21,23^.

#### (3) Dynamic or Slice-Wise Shimming

The OmniShim tool can also isolate sets of slices for “dynamic shimming” or “slice-wise shimming.” In this scenario, separate B_0_ shimming solutions are generated for individual slices or slice groups. This option is particularly useful for techniques using slice-by-slice or true multi-slice acquisitions such as resting-state or task-based functional MRI (fMRI) protocols or multi-slice proton MRSI. By tailoring shim settings to each slice (or group of slices) ^33^, local field inhomogeneities can be corrected more precisely, potentially improving the sensitivity and reliability of applications such as fMRI and MRSI ^29,34–36^. The OmniShim tool can generate shim values for each slice or slice group and store shim values in a format, which the respective scanner can read. However, true dynamic B_0_ shimming needs either additional scanner patches or hardware that allows run time shim current updates. Neither of this is part of this software distribution ^34–38^

In summary, this software tool offers a sophisticated, vendor-agnostic solution for optimizing B_0_ shimming in MRI, combining real shim field calibration and a robustly converging shim algorithm considering real shim field constraints with the practical benefits of flexible ROI selection and seamless cross-vendor applicability.

## METHODS

### Scanners and radiofrequency coils

The herein described data acquisitions have been performed on two distinct human whole body MRI scanners: (i) Siemens Prisma 3T and (ii) Philips dSync 7T. The following radiofrequency coils have been used: (i) 3T Siemens 64-channel receive array and body coil transmission and (ii) 7T Nova Medical 32-ch head coil and 2-channel head only transmission coil.

#### Shim Field Calibration Data Acquisition

Herein, we acquired real shim field calibration data for the B_0_ shim fields generated by the gradient coils, second order, and third-order B_0_ shim coils of the two MRI scanners described in the previous section. This calibration process utilized three-dimensional gradient-recalled echo (3D GRE) sequences with two echo times (TE) with a difference delta TE = 2.3 ms (for 3T system) versus delta TE = 1 ms (for 7T system) between them so that the fat and water was in phase ^29^, The data was collected from a spherical phantom to ensure uniformity and reproducibility. The sequence parameters were as follows: **Philips 7T :** Multi-acquisition (dual echo) with ΔTE = 1.0 ms; TR = 10 ms; flip angle = 20°; Voxel size = 3 × 3 × 3 mm³; 50 slices, FOV = 200 × 200 × 100 mm³, **Siemens 3T:** TE₁ = 4.50 ms, TE₂ = 6.96 ms (ΔTE = 2.46 ms); TR = 651 ms; flip angle = 20°; Voxel size = 3 × 3 × 3 mm³; 60 slices FOV = 192 × 192 × 100 mm³. These data were then processed as described previously to yield scanner specific shim calibration matrices, which were then used for the below described B_0_ shim experiments ^10,11^. For in vivo input B_0_ maps we have used the same scan parameters.

#### Study Design

We have conducted a comprehensive evaluation of our novel B_0_ shimming tool across two distinct magnetic resonance imaging (MRI) scanner platforms, each representing different field strengths and B_0_ shimming capabilities. The experimental setup encompassed: (i) a 3 Tesla (3T) MRI scanner (Siemens Prisma) equipped with second-order shim coils and (ii) a 7 Tesla (7T) MRI scanner (Philips dSync) featuring full second order and third-order shim coils. Our experimental protocol was designed to assess the efficacy of our B_0_ shimming tool in the context of single voxel proton magnetic resonance spectroscopy (MRS), proton magnetic resonance spectroscopic i and functional MRI (fMRI) applications, techniques particularly sensitive to B_0_ field homogeneity._A total of thirty one healthy volunteers (25 male, 6 female, age range : 20 - 30 years) were enrolled in this study and gave informed signed consent in line with the regulations of the local institutional review board (IRB) committee before each scan.

#### 3T MRS and MRSI Data Acquisition

For the 3TSiemens Prisma scanner we conducted in vivo second-order B_0_ shimming with the OmniShim tool in comparison to the vendor provided GRE shim in a cohort of seven subjects (male: five, female: three, age range 20-27 years). The B_0_ shimming performance was evaluated using two distinct spectroscopic techniques: (i) single voxel ^1^H MRS using a vendor provided Point-Resolved Spectroscopy (PRESS) sequence with the following scan parameters: voxel size = 20×20×20 mm³, Echo Time (TE) = 30 ms, Repetition Time (TR) = 2 seconds, Flip Angle = 90°, Voxel angulation = 0° and (ii) ^1^H MRSI using a vendor provided semi-Laser localized chemical shift imaging (CSI) sequence with the following scan parameters: FOV = 95×125×20 mm³, TE = 40 ms, TR = 1.5 seconds, Angulation = 0°, Flip Angle = 90°.

#### 7T MRSI, fMRI and B0 mapping Data Acquisition

At 7T, we extended our investigation to include both second order and third-order B_0_ shimming with the OmniShim tool versus the vendor implemented higher order shim PB volume option, performed on five volunteers (male:3, female: 2 , age range: 20 to 25 years) for ^1^H MRSI and six healthy volunteers (male: 3, female: 2, age range: 20 to 25 years) for fMRI. In addition, we have acquired whole brain B_0_ maps comparing no shimming, first, second and third order PB volume shimming and first, second and third order OmiShim shimming in 5 healthy volunteers (male: three, female: two, age range : 20 to 25 years). The scan protocol comprised: (i) 2D ^1^H FID MRSI^33,44–46,57,58^ with the following scan parameters: TE = 1.21 ms, TR = 320 ms, Flip Angle = 33°, FOV = 200×200 mm, voxel size = 4.4 mm x 4.4 mm, Slice thickness = 12 mm, matrix size = 50×50; (ii) resting state fMRI using a multi-slice single shot EPI sequence covering the whole brain with the following scan parameters: FOV = 220×220×158 mm^3^, TR = 3000 ms, TE = 25 ms; number of dynamics = 120, Flip Angle = 30°, SENSE acceleration factor = 3; multi-band factor = 3 and (iii) Bo maps using a 3D gradient-recalled echo (GRE) sequence with Multi-acquisition (dual echo) with ΔTE = 1.0 ms; TR = 10 ms; flip angle = 20°; Voxel size = 3 × 3 × 3 mm³; 50 slices, FOV = 200 × 200 × 100 mm³.

#### Data Analysis

Single voxel ^1^H MRS data acquired at 3T were exported as raw data and processed with MR-SpectroS^39^ Subsequently these data were fitted with LCmodel^40^ considering the following metabolites: alanine (Ala), aspartate (Asp), creatine (Cr), CSH, γ-aminobutyric acid (GABA), glucose (Glc), glutamine (Gln), glutamate (Glu), glycine (Gly), glutathione (GSH), taurine (Tau), glycerophosphocholine (GPC), lactate (Lac), leucine (Leu), myo-inositol (mI), N-acetylaspartate (NAA), N-acetylaspartylglutamate (NAAG), phosphocreatine (PCr), phosphocholine (PCh), scyllo-inositol (Scyllo), and 2-hydroxyglutarate (2HG) .The respective basis set was simulated using VESPA^41^.Macromolecules were considered in the basis set as described previously^42,43,44^. Proton MRSI data acquired at 3T were processed and fitted with vendor implemented pipelines at the scanner console. Proton spectroscopic imaging datasets acquired at 7T were exported as raw data and reconstructed and processed through established custom written MATLAB and PYTHON pipelines^33,45,46,^. The resulting spectroscopic imaging data were then fitted with LCmodel to extract metabolite information. The respective basis set was simulated using VESPA^41^ Macromolecules were considered in the basis set as described previously. Quantitative corrections and visualization of the metabolite maps was performed as described previously^46^. Functional MRI (fMRI) datasets were preprocessed and analyzed using the CONN toolbox^47, 48^. following the pipeline detailed in the Supplemental Material..

To assess the performance of the OmniShim B_0_ shimming tool, we conducted a comparative analysis against the vendor-provided shimming solutions. This comparison was multifaceted, encompassing: (i) mean and standard-deviation of frequency inside the shim ROI derived fromB_0_ maps acquired through various shim settings; (ii) evaluation of water line width of single voxel MRS data; (iii) evaluation of metabolite spectra from single voxel MRS and MRSI data obtained from various regions of interest within the brain; (iv) assessments of quality of metabolite maps derived from ^1^H MRSI data; (v) Cramer Rao lower bound (CRLB) and (vi) comparing resting state fMRI networks between different shim settings. This comprehensive approach allowed us to quantitatively and qualitatively assess the efficacy of our B_0_ shimming tool in improving B_0_ field homogeneity across different field strengths, B_0_ shimming orders and shim ROIs. Two sample T-tests (unpaired) were performed to verify the statistical significance of the reported results in the figures.

By employing this rigorous methodology, we aimed to demonstrate the versatility and effectiveness of our B_0_ shimming tool in enhancing the quality of spectroscopic and fMRI data acquisition in neuroimaging applications. The inclusion of both 3T and 7T systems in our study provides valuable insights into the tool’s performance across different field strengths and B_0_ shimming systems, potentially offering a significant advancement in the field of MR spectroscopy and imaging.

## RESULTS

### Single-Voxel Spectroscopy at 3T

**Figure 1 (A) and (B)** illustrate single-voxel spectroscopy data collected from the frontal and visual cortices at 3T. When using the OmniShim tool, metabolite spectra show sharper peaks with higher spectral resolution compared to the ones acquired with the vendor shim. In contrast, spectra acquired =with the vendor’s GRE shim exhibit very broad peaks resulting in a largely diminished spectral resolution . This improvement reflects the effectiveness of OmniShim in refining magnetic field homogeneity, ultimately leading to better spectroscopic data quality.

**Figure 1 (C)** further quantifies this enhancement by comparing water linewidths from the same frontal and visual cortex voxels. Box plots reveal a substantial reduction in linewidths with OmniShim B_0_ shimming: the median water linewidth in the frontal cortex decreases from 24.5 Hz (vendor shim) to 9.75 Hz (OmniShim), and in the visual cortex from 19.5 Hz to 9.5 Hz. These findings underscore the pivotal role of improved field homogeneity in producing narrower linewidths, which translate into superior spectral resolution and elevated sensitivity for metabolite detection.

### Spectroscopic Imaging at 3T

**Figure 2 (A) and (B)** present metabolite maps of NAA and tCr acquired with semi-laser ^1^H MRSI at 3T from a slice above corpus callosum, respectively, comparing the data quality obtained with the vendor-provided shim versus OmniShim. Notably, OmniShim consistently yields substantially higher quality metabolite maps with almost complete coverage of the pre-localized ROI highlighting the pronounced advantage of improving the magnetic field’s uniformity.

**Figure2.**
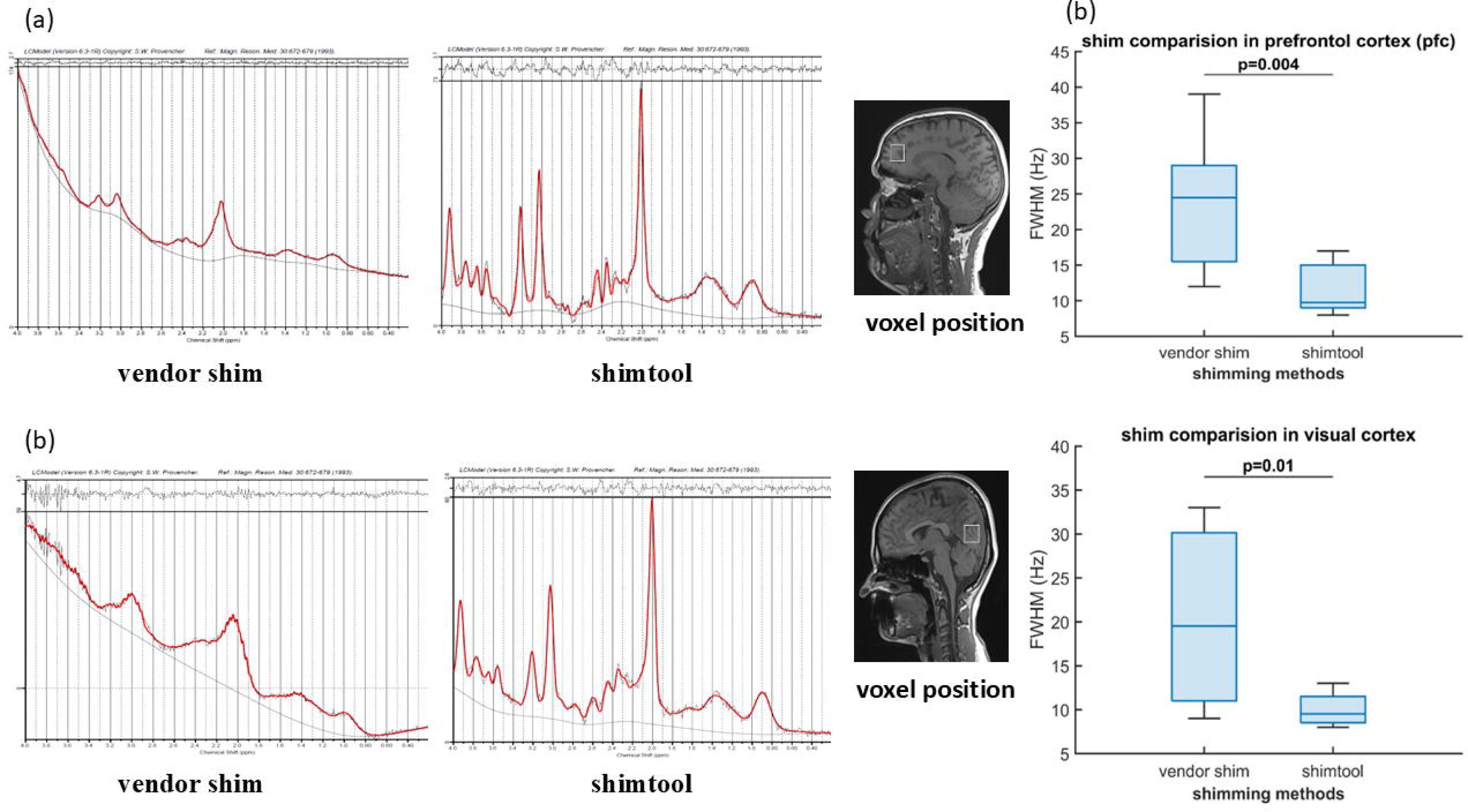
shows comparison of the metabolite maps of N-acetyl aspartate(NAA), and creatine ( Cr ) from the vendor platform. Here we have compared shimming performance of vendor’s default shimming and locally developed shimtool. The order of the shimming is 2^nd^ order.

**Figure 3** presents respective spectra acquired from four voxels spanning different brain regions, illustrating the pronounced impact of OmniShim B_0_ shimming on spectral quality. The improved clarity, sharper peak delineation, and reduced linewidths and reduced lipid contamination observed with OmniShim shimming facilitate more accurate spectral fitting and thus quantification of metabolites and bolster confidence in the resulting spectral measurements.

**Figure3.**
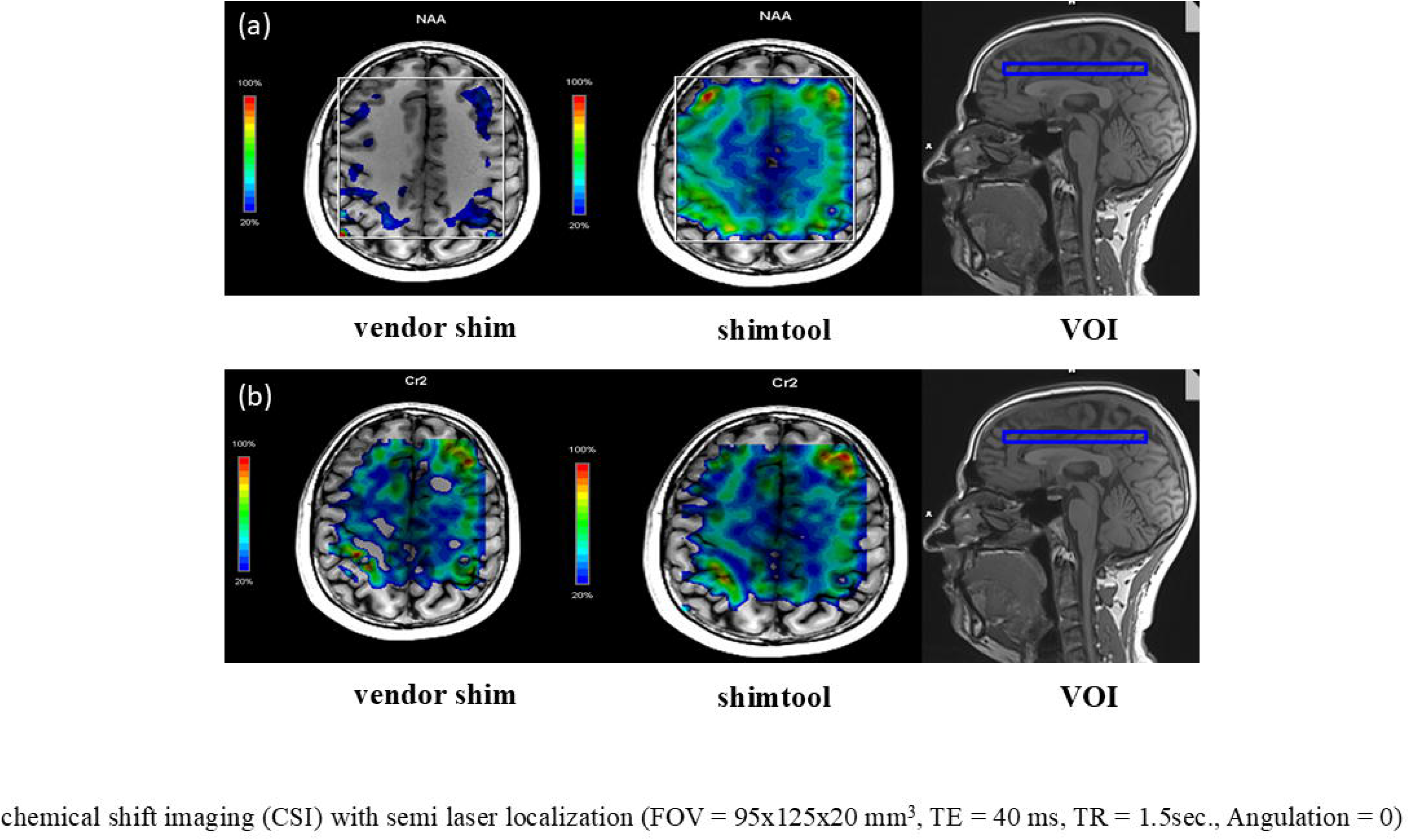
shows the example of metabolite spectra from different voxels. Here we have compared vendor’s default shim v/s our local shim. We can see significant increase in SNR of metabolite spectrum by shimming using locally developed shim. We have used 2^nd^ order shimming to compare both shimming methods. The peak separation between different metabolites is clearly visible when we perform B0 shimming using locally developed shimtool.

### Spectroscopic Imaging at 7T

**Figure 4 (A)** illustrates the planning and voxel placement for the 2D FID MRSI slice positioned to capture a region above the corpus callosum. Figure 4 (B) presents the resultant 7T spectroscopic imaging data, highlighting the substantial improvement in spatial coverage achieved through OmniShim particularly within posterior brain areas.

**Figure4.**
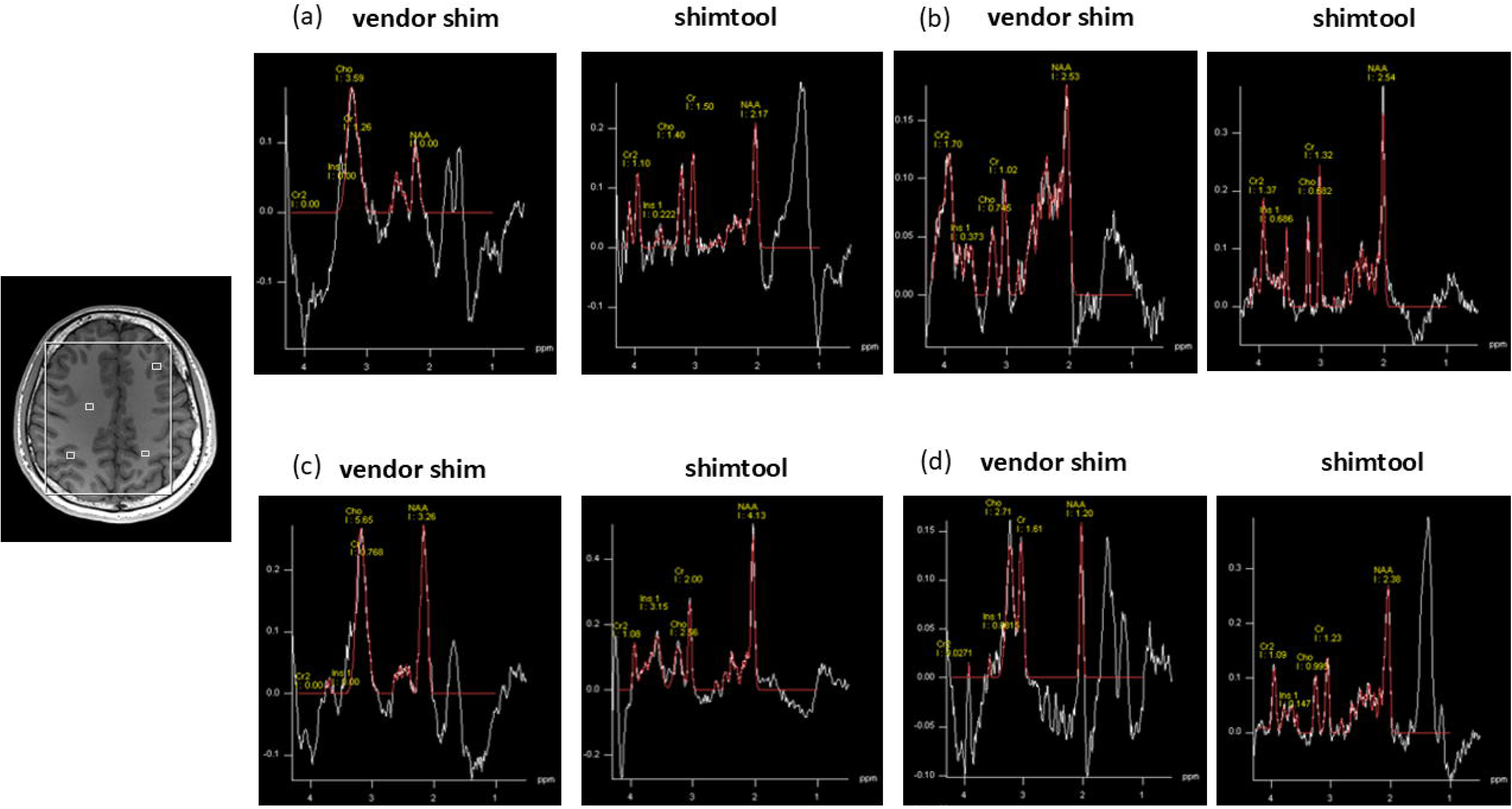
shows the comparison metabolite maps acquired busing 2D FID MRSI at 7T. The first two figures shows the planning of MRSI slice. We have compared vendor implemented 2^nd^ order shim (pencil beam) and compared with our locally developed shimtool (2^nd^ order shim). From the quality of metabolite maps, we can see locally developed shimtool outperformed vendor implemented shimming.

**Figure 5** shows representative spectra from four MRSI voxels across different regions of the brain. We have compared the 2nd order B_0_ shimming of the vendors implemented PB volume shimming and our OmniShim tool at 7T. From the comparison, we can see that spectra acquired through OmniShim shows clearer peak separation between different metabolites compared to metabolite spectra acquired though the vendor implemented PB-volume shim routine with typically detectable brain metabolites being clearly distinguishable. Hence, it OmniShim may deliver more accurate and reproducible MRSI measurements.

**Figure5.**
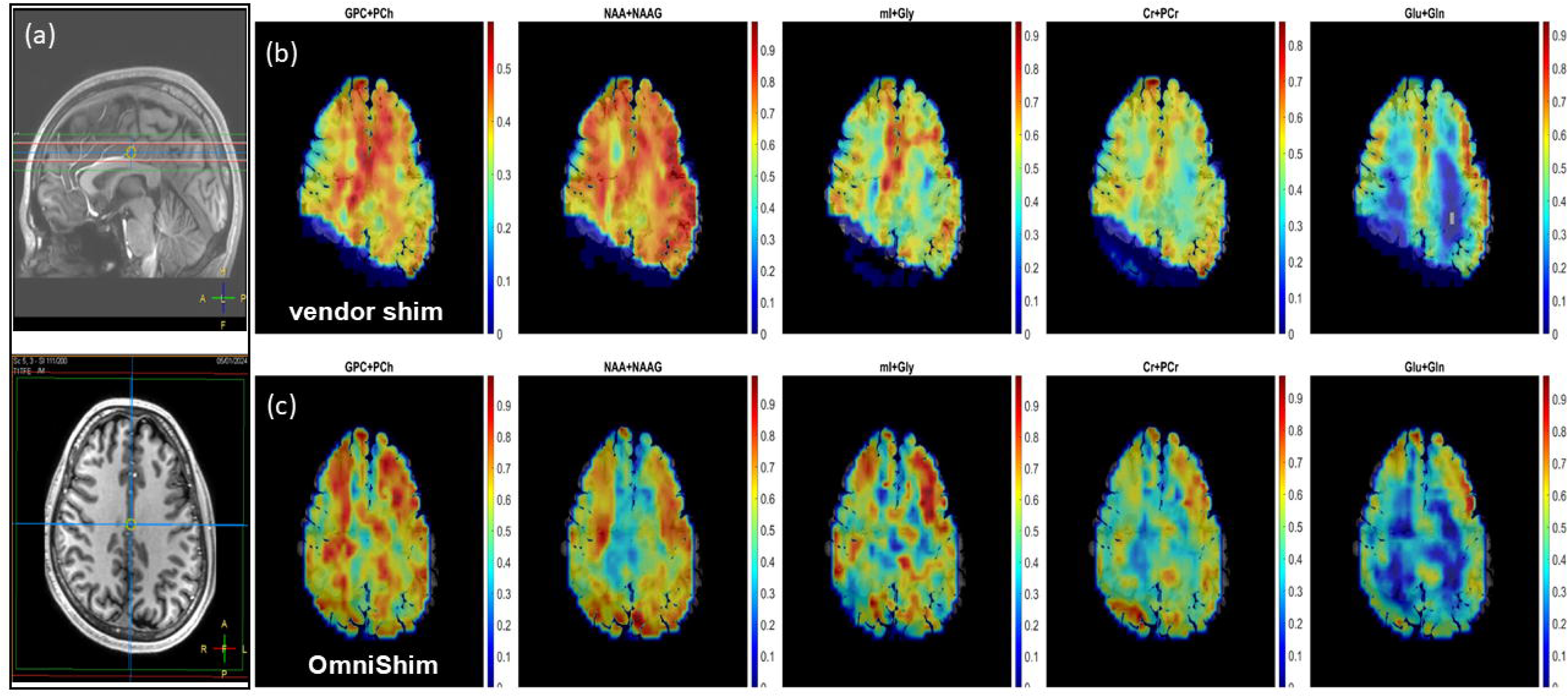
shows the metabolite spectrum from selected voxels across brain. Here we have compared 2^nd^ order vendor implemented shimming (PB volume) with our locally developed shimtool (2^nd^ order shimming). we can see improvement in the quality of the metabolite spectrum by performing shimming with our locally developed shimtool (2^nd^ order shimming). The peak separation between different metabolites is clearly visible when we perform B0 shimming using locally developed shimtool.

**Figure 6** illustrates the comparison of Cramér–Rao lower bound (CRLB) for second-order B_0_ shimming via our vendor-independent OmniShim tool versus the vendor’s built-in PB volume 2^nd^ order shim routine for FID MRSI at 7T. The results demonstrate a pronounced decrease in CRLB after shimming with OmiShim in comparison to PBvolume further underscoring its potential to enhance parameter estimation precision and improve the overall quality of MRSI measurements.

**Figure6.**
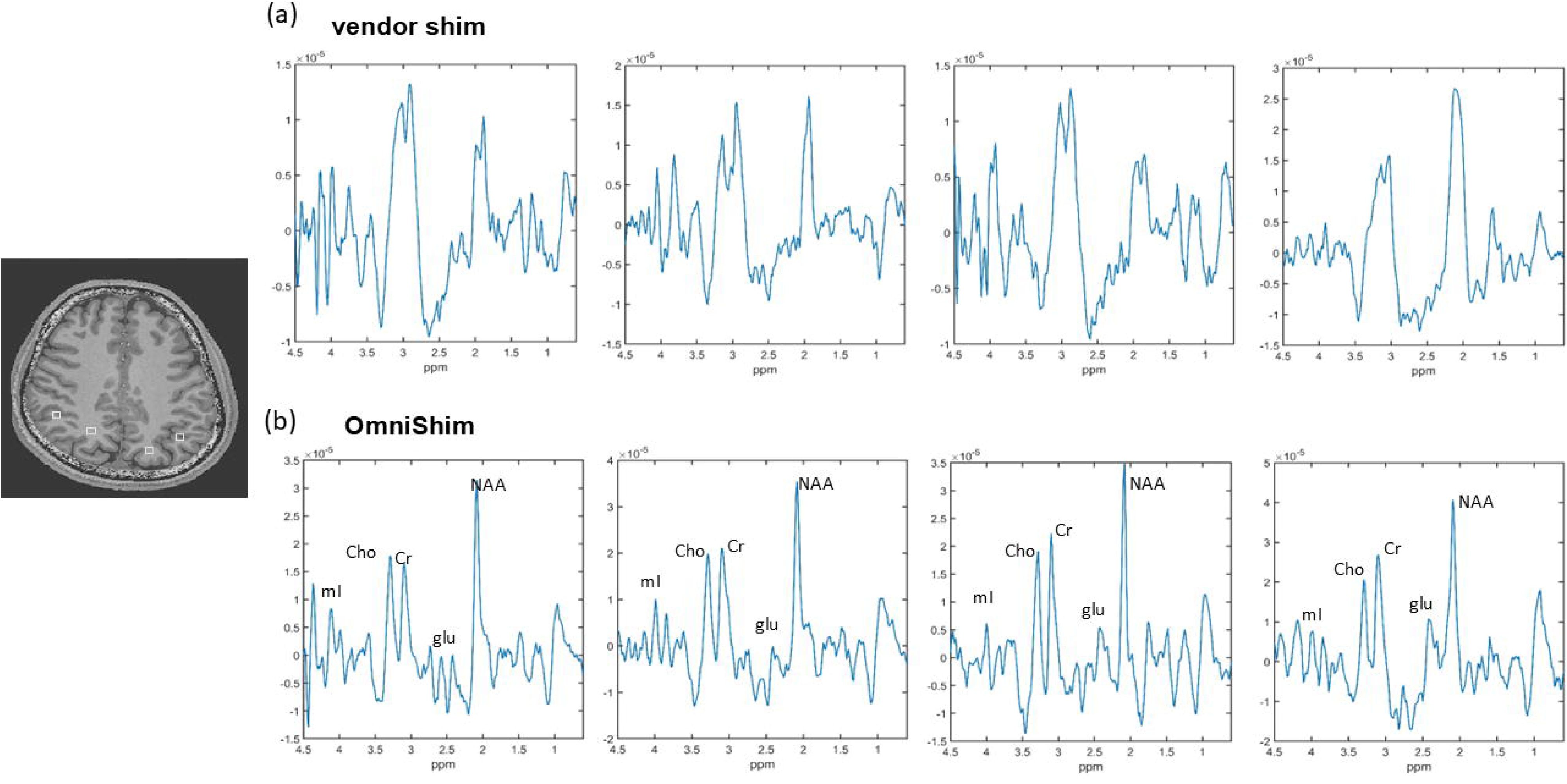
shows Comparison of the Cramér-Rao Lower Bounds (CRLB) for metabolite quantification between vendor-implemented B₀ shimming and the locally developed shim tool. the fitting uncertainty in metabolite concentration maps is significantly reduced, reflecting more reliable spectral fitting and quantification.

### B0 mapping at 7T

**Figure 7** present multi-slice B_0_ maps, along with their corresponding standard deviations, for different shimming approaches: (i) PB volume higher order shimming (vendor), (ii) autoshim first order shimming (vendor) and (iii) OmniShim for first, second and third order shimming in slices located below the corpus callosum at 7T. It is demonstrated that OmniShim provides the better compromise between different regions in presence of the strong local inhomogeneity in the frontal cortex, while the PB volume shimming focusses on reducing the frontal cortex inhomogeneity at the expense of worse shimming everywhere else including the occipital lobe. This is caused by the incomplete sparse information about spatial inhomogeneity that the projections used for PBvolume shimming provide. A related analysis in brain regions above the corpus callosum (Supplemental Figure 7) shows minimal difference between the first- and second-order shimmed B_0_ maps acquired with both the vendor PB volume shim and the OmniShim, primarily because the magnetic field can be readily homogenized due to the lack of strong local inhomogeneity. Consequently, second-order shimming confers little additional benefit in this specific location, highlighting that higher-order B_0_ shimming strategies are most advantageous in more challenging anatomical regions suffering from greater susceptibility-induced inhomogeneities.

**Figure7.**
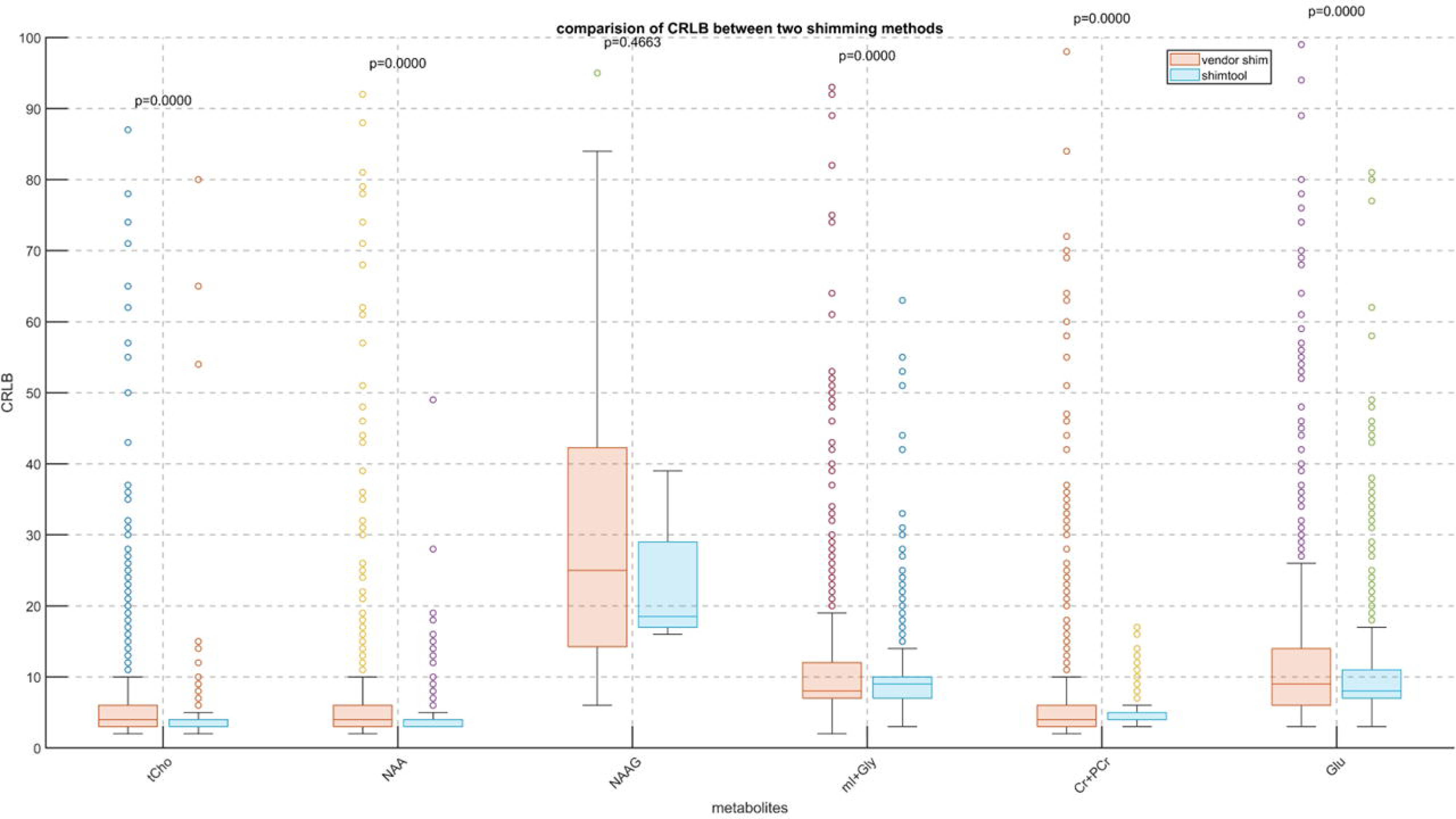
shows comparison of B_0_ maps acquired without shimming B_0_ maps and using different shim orders with vendor’s implemented shim routine as well as our locally developed shimtool. Here, we are trying to shim few B_0_ map slices below corpus collosum. The results shows comparison of B0 maps with various shim orders. We can see B0 maps acquired using vendors implemented third order shimming is very bad.

**Figure 8** shows respective box plots confirming the better allover performance of OmniShim versus the vendor provided PBvolume and Autoshim options for brain regions below the corpus callosum. Altogether 2^nd^ order OmniShim with first order PB volume shimmed input maps (option 9) yield the lowest standard deviation across the entire shim ROI.

**Figure8.**
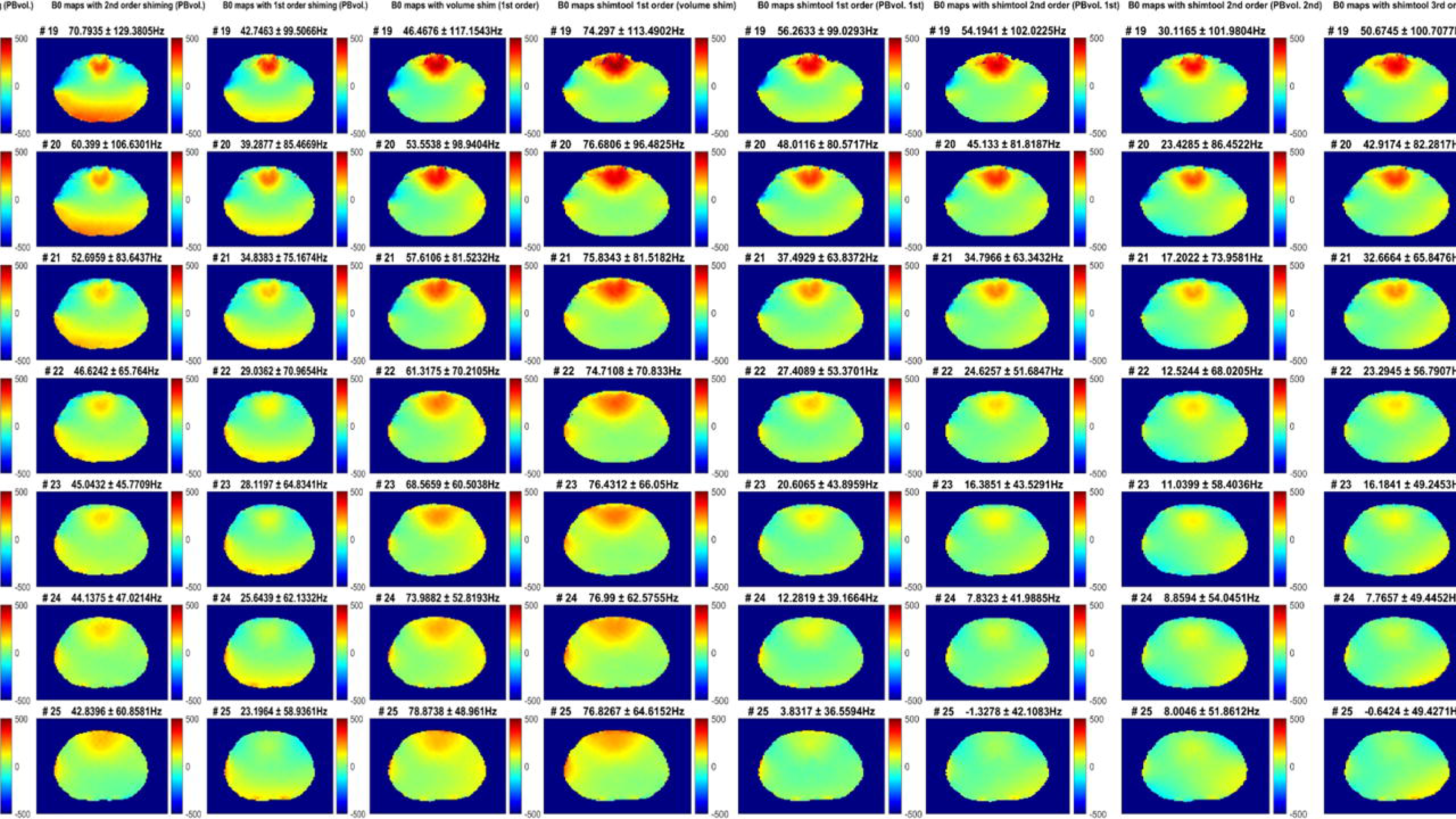
shows comparison of standard deviation of shimmed/Unshimmed B0 map slices below corpus callosum using different shimorder (1^st^ to 3^rd^ order). We have compared shimming performance of vendor implemented shimming routine and our locally developed shimtool. Also, we have used different B0 maps using different shimorder as an input in our tool and measured performance of our shimtool with different shimmed B0 maps as an input.

### Resting-State fMRI at 7T

**Figure 9** presents a few representative slices showing multi-slice single shot EPI data from resting-state fMRI acquisitions after applying the default geometric distortion correction implemented at the scanner console at 7T. We have compared 2^nd^ order shimming performance of the vendor’s PBvolume shimming routine and our OmniShim shimtool. The results demonstrate that the OmniShim shimming method minimizes signal dropouts in a lot of different brain regions temporal lobes, cerebellum, brain stem, ear channels and frontal cortex compared to the vendor’s shimming solution, underscoring the advantages of OmniShim based B_0_ field optimization in fMRI applications.

**Figure9.**
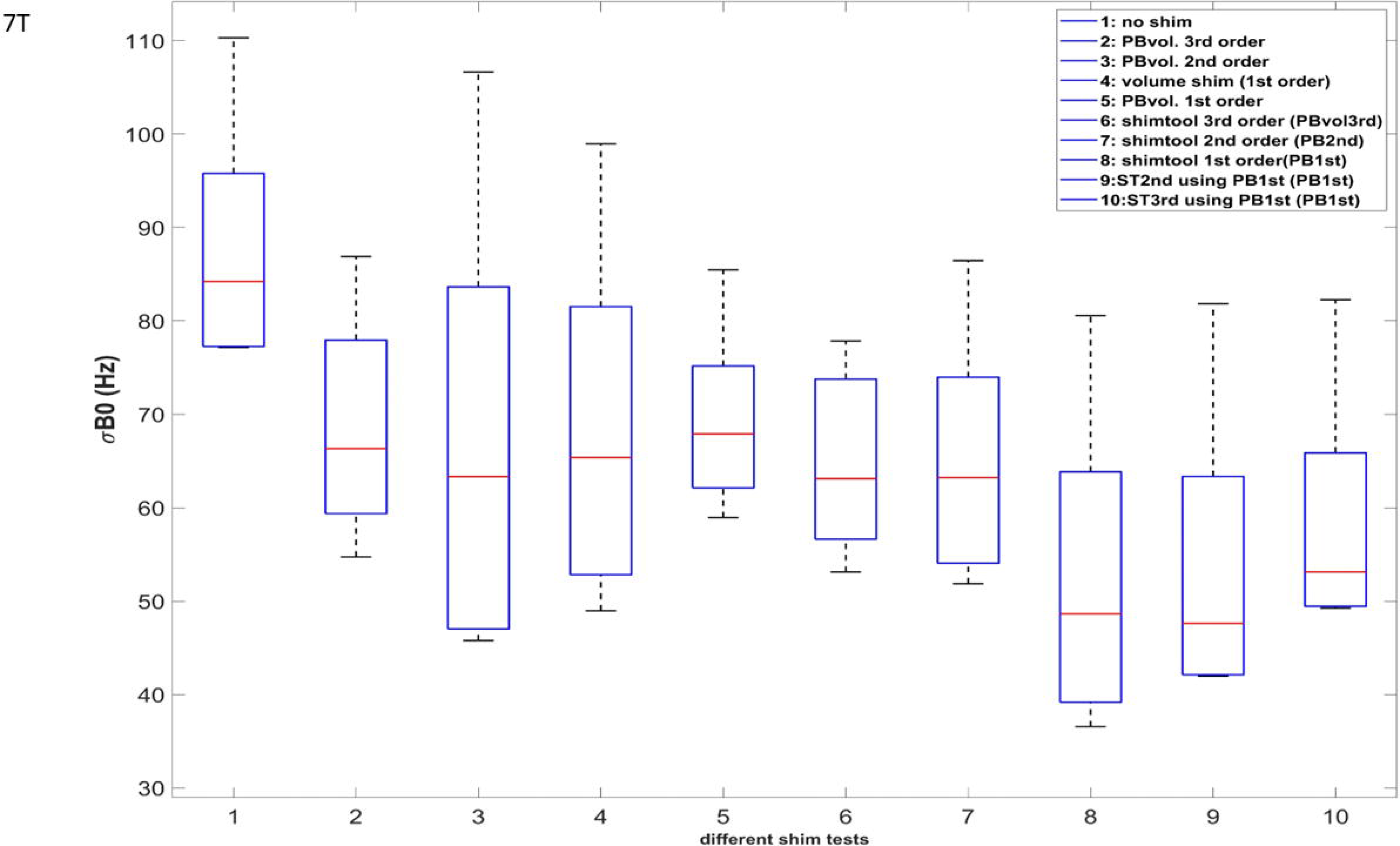
shows 2^nd^ order shimming comparison of vendor implemented shim routine and our shimtool in the resting state fMRI application. We have compared performance of 2^nd^ order shimming vendors implemented the shimming routine and our locally developed shimtool. The comparison is marked by a rad arrow in all the figures. From the image quality we can see the locally developed shimtool improve image quality compared to vendors shimming.

**Figure 10** illustrates functional connectivity network maps in the posterior cingulate cortex (PCC; MNI: 1, –61, 38), the posterior cerebellar lobe (Crus I/II; MNI:0, -63, -30), and the anterior cerebellar lobe (Crus I/II ; MNI: 0, -79, -32). We have compared 2^nd^ order shimming with the vendor implemented PBvolume shim routine with the OmniShim shimtool. For the PCC seed region, Vendor shim exhibits the lack of detectable functional connectivity to the medial PFC and the bilateral angular gyri of the default mode network (DMN). In contrast, OmniShim shimming yields more reliable DMN mapping showing all components of this network. In the posterior cerebellum OmniShim data show clear functional connectivity between cerebellum Crus I and II on the left and right side, while the vendors shim lacks some of parts of this network. Anterior cerebellar rs-fMRI data show clear connectivity inside the right Crus I using OmniShim, while we cannot see any such activation using vendor PBvolume shimming.

**Figure10.**
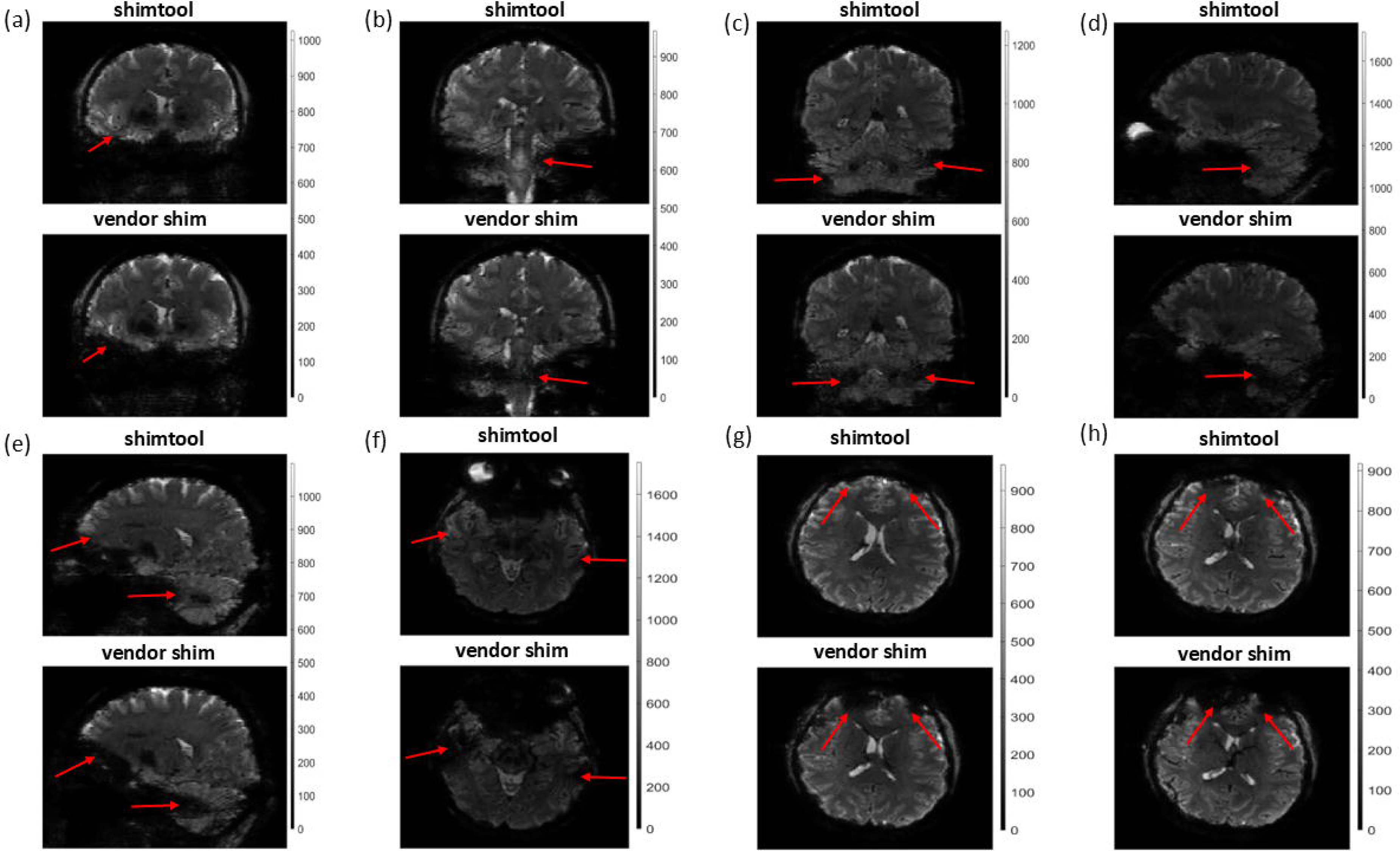
shows comparison of different connectivity networks in PCC Cerebellar Posterior and Cerebellar Anterior regions. The blue arrow shows connectivity network across the regions. here we have compared 2^nd^ order shimming performance between vendors implemented shim routine v/s our locally developed shimtool.

This comparison demonstrates that OmniShim leads to superior functional connectivity results across all three networks implying more reliable resting state fMRI at 7T. These findings emphasize the potential of OmniShim to refine our understanding of neural circuit dynamics, particularly in subcortical and cerebellar structures critical for motor coordination and cognitive processing.

Altogether, the OmniShim shimming method can be employed to consistently optimize the B_0_ homogeneity in different ROIs raging from spectroscopy voxels to large brain volumes, leading to enhanced spectroscopy and image quality and reduced sensitivity to B_0_ inhomogeneities.

## Discussions

Magnetic field (B_0_) inhomogeneity remains a persistent challenge in magnetic resonance imaging (MRI) and spectroscopy (MRS) at 3T and especially at ultrahigh fields such as 7 T. When the B_0_ field is nonuniform, image artifacts emerge, signal-to-noise ratio (SNR) degrades, signal dropouts occur, and the accuracy of quantitative measurements deteriorates. Additionally, these inhomogeneities can lead to broader resonance lines in MRS, complicating the detection and quantification of metabolic peaks. Consequently, the design and optimization of a robust B_0_ shimming strategy are critical for maintaining high-quality data and reliable interpretations.

In practice, it is essential to characterize the real shim fields of each specific shim system of an MRI scanner prior to implementing a B_0_ shimming protocol. Although many scanner manufacturers provide built-in B_0_ shimming hardware and software, these vendor-specific solutions can vary substantially in performance and flexibility and typically don’t consider real shim fields. This variability, coupled with the increasing complexity of higher-order shim systems with shim coils that often exhibit shim fields that do not match ideal spherical harmonic profiles^11^ highlight the need for a vendor-independent B_0_ shimming tool that considers real shim fields and can be calibrated once and then applied consistently across different scans and applications. However, the real shim field calibration process is only helpful if the impurity in the shim coils designed to produce certain shim fields are moderate. With very larger impurities such as the very strong linear cross terms in 3^rd^ order shim coils in whole body 7T MRI scanners ^11,34^ higher order B_0_ shimming cannot live up to its potential^10,20^ and are rather ineffective even after the real field calibration process. In addition to real shim field calibration, a shim algorithm accounting for shim field constraints^21,25^ and robustly converging in presence of noise or localized artefacts in the input Bo maps in combination with flexible ROI definition options were shown to be advantageous and provided by the herein described OmniShim tool.

Figures 2 through 10 provide a comprehensive comparison between the standard Siemens and Philips supplied B_0_ shimming procedures and the OmniShim B_0_ shimming method applied to various brain regions, shim ROIs and acquisition conditions at 3T and 7T. The analysis spans single-voxel spectroscopy in the frontal and visual cortices, spectroscopic imaging above and below the corpus callosum, as well as resting-state functional MRI (fMRI). Overall, the OmniShim shimming approach demonstrates considerable improvements in both spectroscopic and functional imaging quality, as evidenced by narrower linewidths, better discernable spectral peaks and a reduction in artefacts in MRS and MRSI and a reduction of signal dropouts along with a better delineation of known functional networks in functional imaging data.

Localized shimming reduces B₀ inhomogeneity improving peak separation of key metabolite resonances (e.g., NAA, Cr, Cho, Glu and mI). This sharpening of spectral lines minimizes peak overlap of adjacent resonances such as choline and creatine allowing unambiguous peak separation. It also prevents lipid contamination from skull lipid in brain MRSI, which negatively impacts the quantification of metabolite peaks adjacent to the main lipid resonance such as=N-acetylaspartate. Moreover, water suppression relies on frequency selective saturation and thus on high B_0_ homogeneity across the field of view. Altogether, the underlying baseline becomes flatter in presence of superior B_0_ shimming and thus water suppression, reducing spectral fitting errors. The combination of linewidth narrowing, amplitude gain along with artefact control translates into a net SNR increase across metabolite peaks in MRS and MRSI and a much-improved spatial coverage in MRSI. Comparison of the Cramér-Rao Lower Bounds (CRLB) for metabolite quantification between vendor-implemented B_0_ shimming and the OmniShim tool demonstrate that the latter yields lower CRLB values, indicating improved accuracy in metabolite estimation. Such gains in spectral quantification accuracy enhance the reliability of metabolic biomarkers, particularly in regions with strong local magnetic susceptibility gradients (e.g., frontal lobes, near sinuses). In practice, this can improve the detection of low-concentration metabolites (e.g., myo-inositol, glutathione) and sharpen the delineation of pathological versus healthy tissue. In case of fMRI OmniShim yielded a marked reduction in signal loss near areas with strong local B_0_ inhomogeneity including the frontal cortex, brain tissue near the ear channels, the temporal lobes, cerebellum and brain stem. This in turn yielded more robust functional connectivities in respective brain networks. The PCC connectivity reflects the DMN and hence introspective and memory-related processes. Posterior cerebellar connectivity shows Crus I/II map highlighting cerebellar contributions to higher-order cognition and executive function. Anterior cerebellar connectivity shows Crus I/II map highlighting cerebellum’s role in sensorimotor coordination. Our OmniShim tool has increased suprathreshold cluster volume, demonstrating that enhanced B₀ homogeneity directly translates into stronger, more spatially extensive functional networks. These functional connectivity results underscore how sub-optimal B_0_ shimming can bias resting-state connectivity—particularly around air-tissue interfaces—and how our shimtool mitigates these effects, yielding more faithful maps of both default-mode and cerebellar networks.

The B_0_ mapping experiments demonstrated the positive impact of a properly calibrated higher order B_0_ shim system in the lower brain below the corpus callosum that exhibits most non-spherical air-tissue interfaces and hence local inhomogeneity. Due to the poor design of the third order shim systems at 7T whole-body human MRI scanners with strong linear contaminations^11^ second order B_0_ shimming showed the best performance.

Compared to projection-based shimming techniques (e.g., FASTMAP and its derivatives)^12,13,49,50^ our vendor-independent OmniShim tool explicitly accounts for all sources of inhomogeneity in a 3D shim ROI. As demonstrated herein this is especially important in case of complex B_0_ inhomogeneities such as in the lower brain. While projection-based B_0_ shimming was recommended for single voxel MRS by a recent consensus paper, image-based shimming is generally recommended for MRSI and fMRI^51^. In addition, none of the published projection-based methods consider real shim fields. One early study has investigated as to how much improvement can be achieved using B_0_ higher-order’s shims compared with the linear shims alone using projection mapping methods in chemical shift imaging (CSI)^16^ More recently, the advantages of higher order shim systems have been shown with image based B_0_ shim approaches clearly^10,20,38^. OmniShim is an image-based shim method considering hardware constraints with respect to maximum shim currents and shim field sensitivities of the built-in shim coils along with residual field distortions introduced by impure higher-order shim coils. Following a one-time calibration of the scanner specific real shim fields, this approach can be especially advantageous for larger or irregularly shaped regions of interest (ROIs), such as those typically encountered in magnetic resonance spectroscopic imaging (MRSI) and non-localized MRS studies, fMRI, spinal cord MRI, cardiac and body MRI as well as musculoskeletal MRI.

While several toolboxes are available for B0 shimming using vendor-provided shim hardware^8,21,35^, and others have demonstrated the use of external shim systems employing either full spherical harmonic basis sets^19^ or custom non-spherical harmonic coil configurations^52–54^, to the best of our knowledge, there is currently no open-source, all-in-one software platform that supports a wide range of B0 shimming strategies and operates across different MRI vendors. To address this gap, we developed and validated OmniShim, a vendor-independent B0 shimming tool compatible with multiple platforms. OmniShim was tested across different field strengths (3T and 7T) and vendors (Siemens and Philips). Validation experiments included both healthy volunteers and brain tumor patients, demonstrating the tool’s flexibility and robustness in diverse shimming scenarios.

## Limitation

This vendor-independent OmniShim shimming tool fully leverages the potential of existing B_0_ shim coils provided by the scanner manufacturer. However, we previously showed that vendor-supplied higher-order shim coils do not produce purely spherical harmonic fields, with the third-order shim coils in 7T human MRI scanners exhibiting particularly notable deviations ^11^. As a result, in practical scenarios, second-order shim solutions tend to yield the most consistent outcomes in spite of real field calibrations. While OmniShim is theoretically capable of generating shim solutions up to sixth order, the vendor system examined herein do not include coils beyond the third order, and their non-ideal performance limits its benefits. Despite these imperfections of current 3^rd^ order shim systems higher order B_0_ shim coils remain highly promising solutions for mitigating field inhomogeneities in challenging anatomical regions, such as the sinuses., The presented OmniShim shimming framework is readily adaptable to any future improved vendor implemented shim hardware as well as external shimming hardware designed with pure spherical harmonic fields up to the full 4^th^ order^10,20,38^ or custom non-spherical harmonic implementations ^37,53–56^ , offering broad flexibility.

Furthermore it should be noted, that for smaller ROIs such as MRS voxels, the limited number of acquired field map points may compromise the effectiveness of the shim solution. Although it is generally recommended to acquire higher-resolution B_0_ maps in these cases, there is an inherent tradeoff: using a higher-resolution field map can better capture local inhomogeneities but will also increase acquisition time and complexity. Striking the right balance between capturing sufficient detail within the VOI and maintaining practical scan durations remains a central challenge. By systematically calibrating and leveraging our vendor-independent shim tool, researchers and clinicians can mitigate B_0_ inhomogeneity more effectively, thereby improving both image quality and the accuracy of quantitative MRS-based measurements.

## Conclusion

In summary, the data presented in these figures underscore the benefits of OmniShim in improving the quality of both spectroscopic and functional MRI data. By enhancing field homogeneity, OmniShim results in higher SNR, sharper linewidths and thus higher spectral resolution artifact reduction and thus higher quantification accuracy and improved spatial coverage in MRS and MRSI. IN addition, more accurate functional connectivity maps, particularly in regions like the cerebellum and frontal cortex can be derived. These improvements have the potential to significantly enhance the sensitivity and precision of functional and metabolic neuroimaging, enabling more reliable brain mapping and better understanding of brain function and connectivity.

Access can be granted upon reasonable inquiry, and distribution is intended to support academic and scientific use.

## Acknowledgement

This work was performed at Advance Imaging Research Center at University of Texas Southwestern Medical center Dallas. This work was supported by Cancer Prevention and Research Institute of Texas (CPRIT) grant / RR180056.

## Processing steps to Analyze resting state fMRI data

### Methods

Results included in this manuscript come from analyses performed using CONN[1] (RRID:SCR_009550) release 20.a[2] and SPM[3] (RRID:SCR_007037) release 12.7771.

#### Preprocessing

Functional and anatomical data were preprocessed using a flexible preprocessing pipeline[4] including realignment with correction of susceptibility distortion interactions, slice timing correction, outlier detection, direct segmentation and MNI-space normalization, and smoothing. Functional data were realigned using SPM realign & unwarp procedure[5], where all scans were coregistered to a reference image (first scan of the first session) using a least squares approach and a 6 parameter (rigid body) transformation[6], and resampled using b-spline interpolation. Temporal misalignment between different slices of the functional data (acquired in interleaved Siemens order) was corrected following SPM slicetiming correction (STC) procedure[7,8], using sinc temporal interpolation to resample each slice BOLD timeseries to a common mid-acquisition time. Potential outlier scans were identified using ART[9] as acquisitions with framewise displacement above 0.9 mm or global BOLD signal changes above 5 standard deviations[10,11], and a reference BOLD image was computed for each subject by averaging all scans excluding outliers. Functional and anatomical data were normalized into standard MNI space, segmented into grey matter, white matter, and CSF tissue classes, and resampled to 2 mm isotropic voxels following a direct normalization procedure[11,12] using SPM unified segmentation and normalization algorithm[13,14] with the default IXI-549 tissue probability map template. Last, functional data were smoothed using spatial convolution with a Gaussian kernel of 8 mm full width half maximum (FWHM).

#### Denoising

In addition, functional data were denoised using a standard denoising pipeline[15] including the regression of potential confounding effects characterized by white matter timeseries (10 CompCor noise components), CSF timeseries (5 CompCor noise components), motion parameters and their first order derivatives (12 factors)[16], outlier scans (below 48 factors)[10], session and task effects and their first order derivatives (8 factors), and linear trends (2 factors) within each functional run, followed by bandpass frequency filtering of the BOLD timeseries[17] between 0.01 Hz and 0.1 Hz. CompCor[18,19] noise components within white matter and CSF were estimated by computing the average BOLD signal as well as the largest principal components orthogonal to the BOLD average, motion parameters, and outlier scans within each subject’s eroded segmentation masks. From the number of noise terms included in this denoising strategy, the effective degrees of freedom of the BOLD signal after denoising were estimated to range from 137.5 to 169.9 (average 160.5) across all subjects[11].

#### First-level analysis

SBC_01: Seed-based connectivity maps (SBC) and ROI-to-ROI connectivity matrices (RRC) were estimated characterizing the patterns of functional connectivity with 32 HPC-ICA network ROIs[2]. Functional connectivity strength was represented by Fisher-transformed bivariate correlation coefficients from a weighted general linear model (weighted-GLM[20]), defined separately for each pair of seed and target areas, modeling the association between their BOLD signal timeseries. Individual scans were weighted by a boxcar signal characterizing each individual task or experimental condition convolved with an SPM canonical hemodynamic response function and rectified.

#### First-level analysis

ICC_01: Intrinsic Connectivity maps (IC) characterizing network centrality at each voxel were estimated as the root mean square (RMS) of all short- and long-range connections between a voxel and the rest of the brain[21]. Connections were computed from the matrix of bivariate correlation coefficients between the BOLD timeseries from each pair of voxels, estimated using a singular value decomposition of the z-score normalized BOLD signal (subject-level SVD) with 64 components separately for each subject and condition[1]. Last, IC measures across voxels were rank sorted and normalized separately for each individual subject and condition using a Gaussian inverse cumulative distribution function with zero mean and unit variance.

#### Group-level analyses

were performed using a General Linear Model (GLM[22]). For each individual voxel a separate GLM was estimated, with first-level connectivity measures at this voxel as dependent variables (one independent sample per subject and one measurement per task or experimental condition, if applicable), and groups or other subject-level identifiers as independent variables. Voxel-level hypotheses were evaluated using multivariate parametric statistics with random-effects across subjects and sample covariance estimation across multiple measurements. Inferences were performed at the level of individual clusters (groups of contiguous voxels). Cluster-level inferences were based on parametric statistics from Gaussian Random Field theory[23,24]. Results were thresholded using a combination of a cluster-forming p< 0.001 voxel-level threshold, and a familywise corrected p-FDR < 0.05 cluster-size threshold[25].

