## Supplemental material for "OmniShim - a vendor-independent B0 Shimming software toolbox"

### SUPPORTING INFORMATION

**Figure. S1.** This figure shows the layout of the shimtool user interface. This user interface shows the different methods of shimming and options for ROI drawings. We can map mask in any plane. The tool required dicom image as an input B0 map.

**Figure. S2.** This figure shows the single voxel and MRSI masks in coronal plane in shimtool interface.

**Figure. S3.** The figure illustrates how our shimtool generates and deploys randomly placed regions of interest (ROIs) within a given volume to assess $B_0$ homogeneity and shim performance.

**Figure. S4.** This figure shows the predictions of corrected $B_0$ maps after calculating shims and applying shims to original $B_0$ maps to check shim performance.

**Figure. S5.** This figure shows the concept of ROLI algorithm and how shimtool will select region of less interests. Tool can calculate weights using Region of less interests (ROLI) and try to shim regions causing inhomogenities in the Region of interest.

**Figure S6.** present multi-slice $B_0$ maps in slices located above the corpus acquired without shim and with different shim orders using vendors implemented shimming (PBvolume) and locally developed shimtool at 7T.

**Figure S7.** presents multi-slice B0 maps, along with their corresponding standard deviations, for different shimming approaches (first order versus second order) in slices located below above the corpus callosum at 7T. The data indicate minimal discrepancies between the first- and second-order shimmed B0 maps in this region, primarily because the magnetic field can be readily homogenized in the relatively simple anatomical environment above the corpus callosum. Consequently, second-order shimming confers little additional benefit in this specific location, highlighting that higher-order B0 shimming strategies are most advantageous in more challenging anatomical regions characterized by greater susceptibility-induced inhomogeneities.

Supplement figure 1:

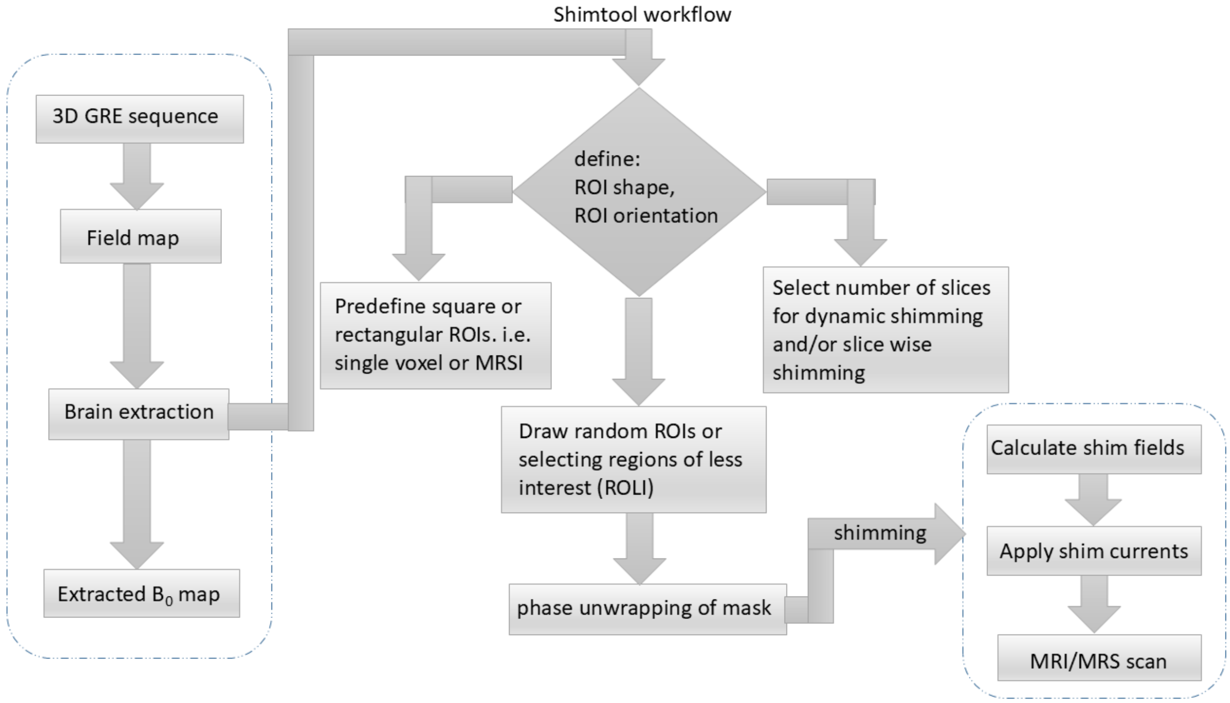


Supplement figure 2:

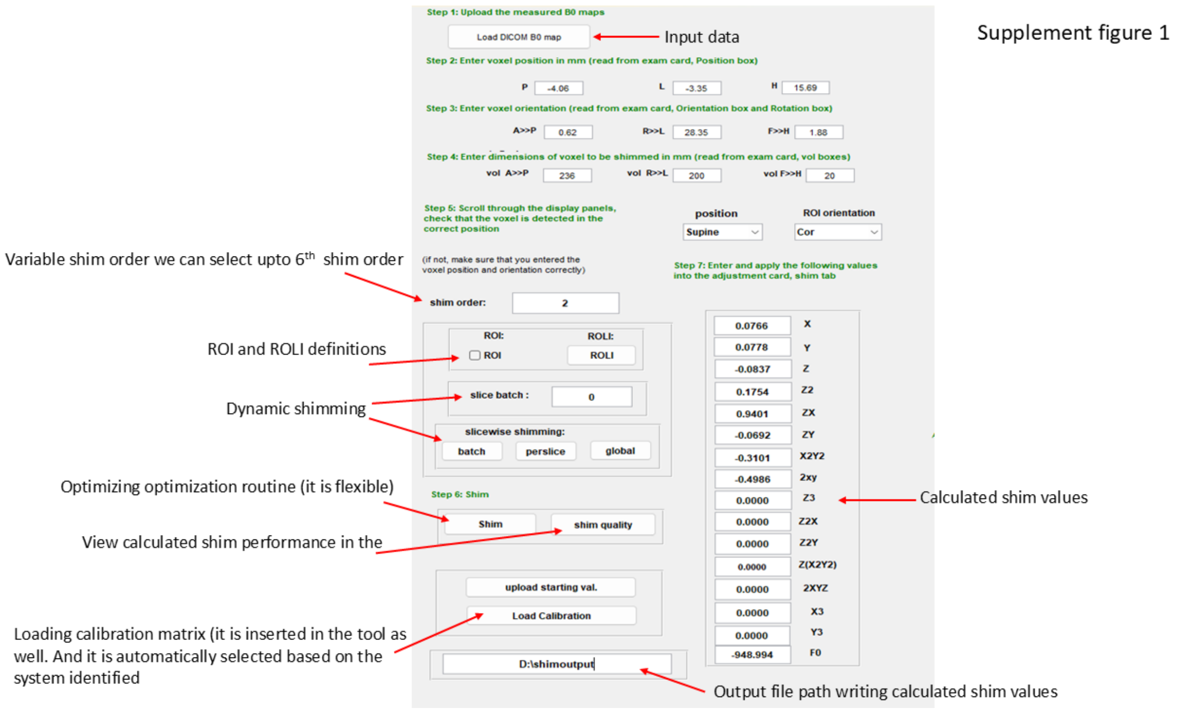

Supplement figure 3:

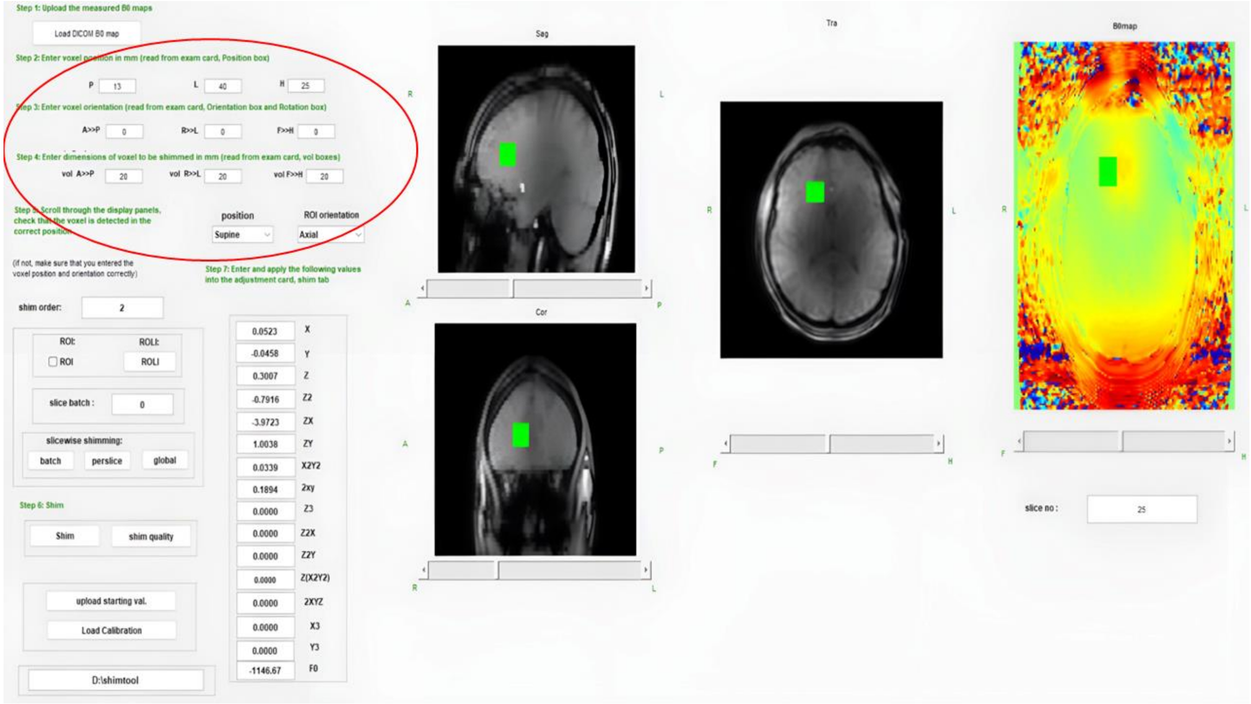


Supplement figure 4:

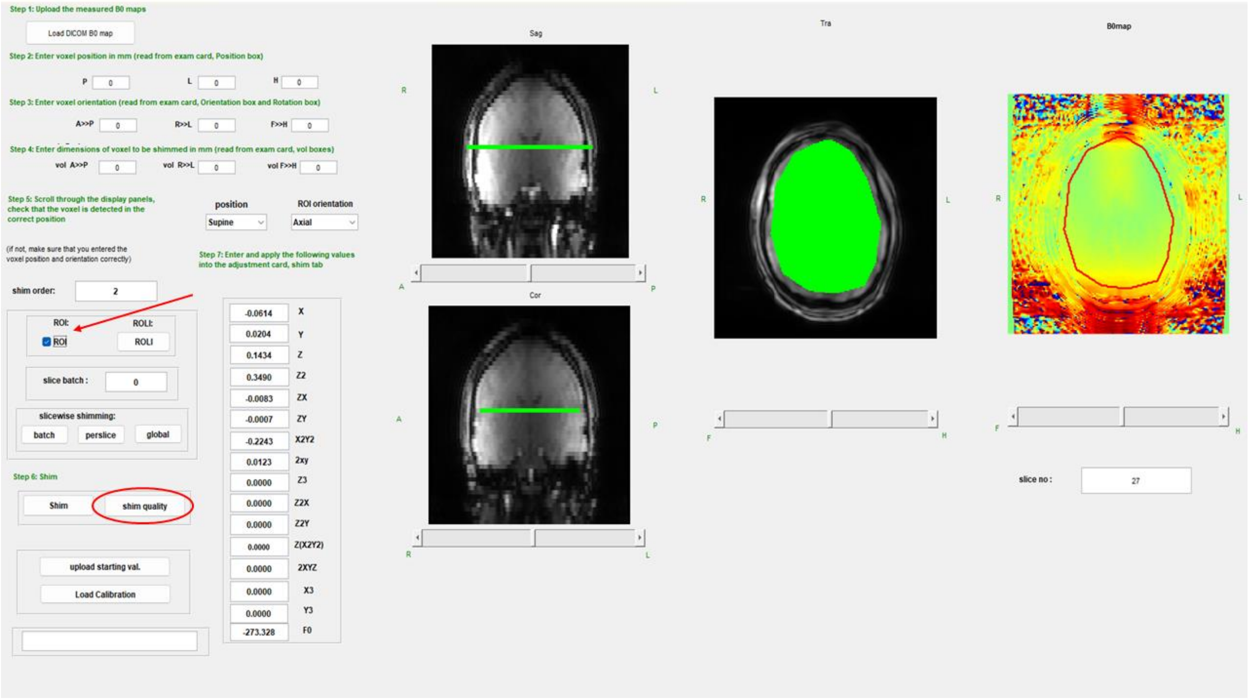

Supplement figure 5:

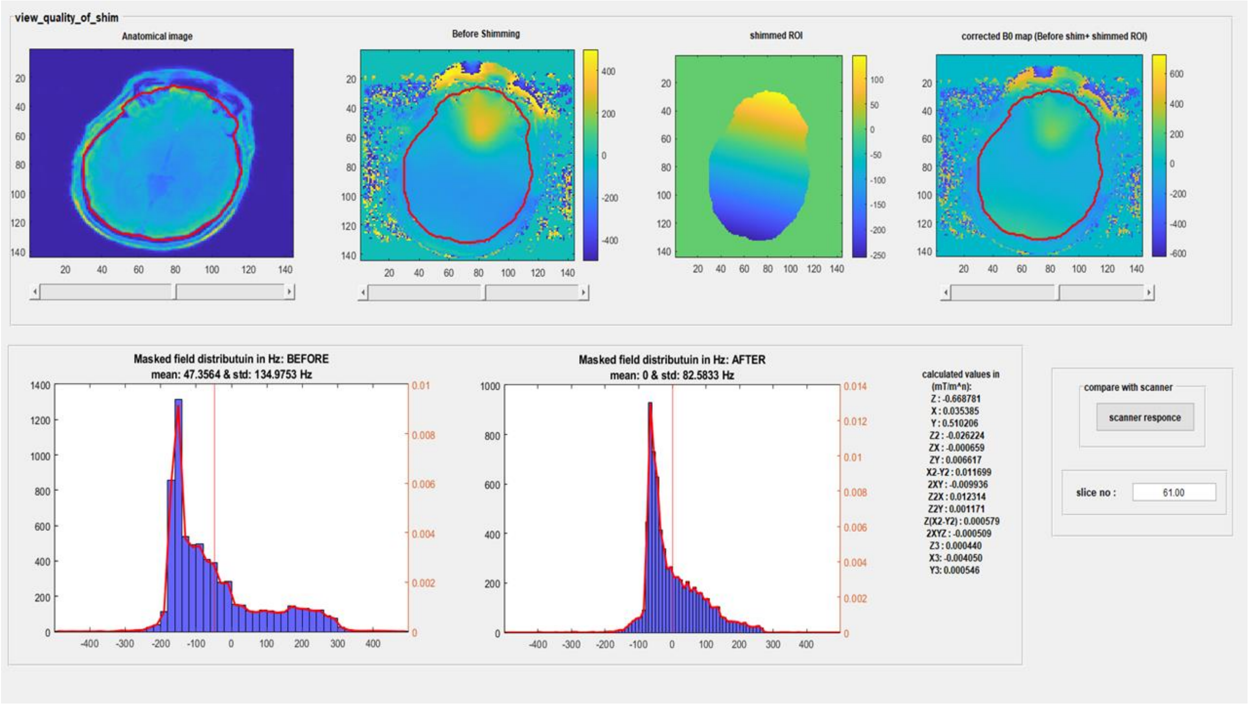

Supplement figure 6:

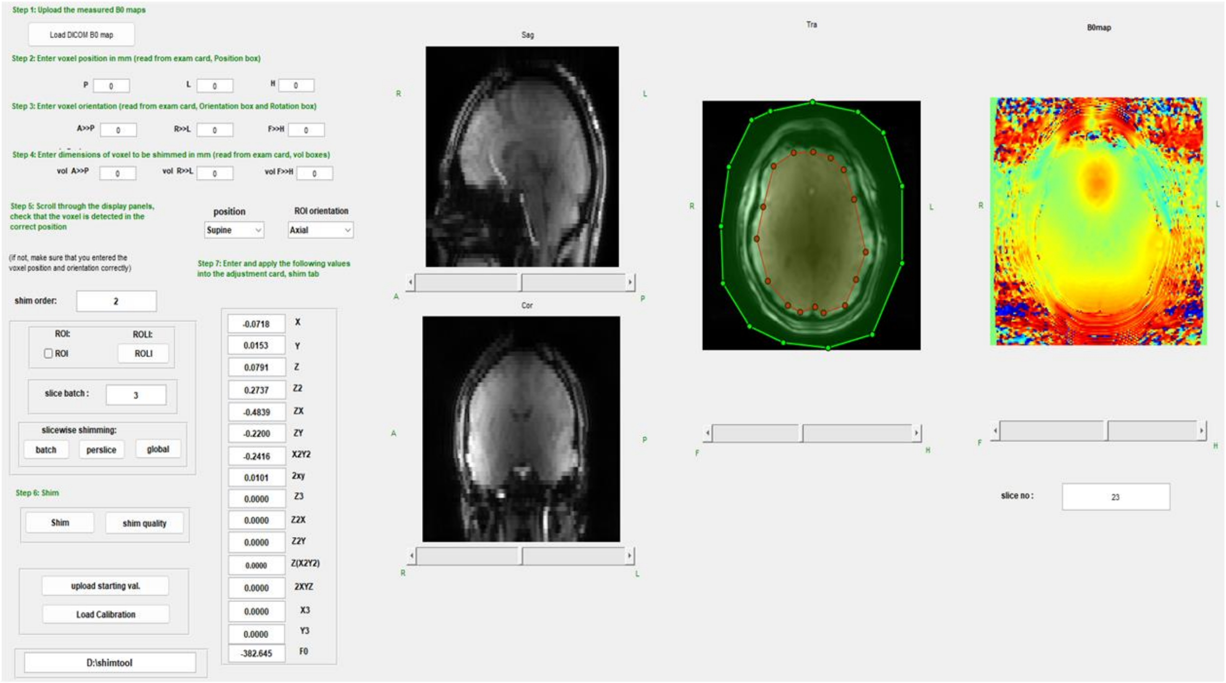

Supplement figure 7:

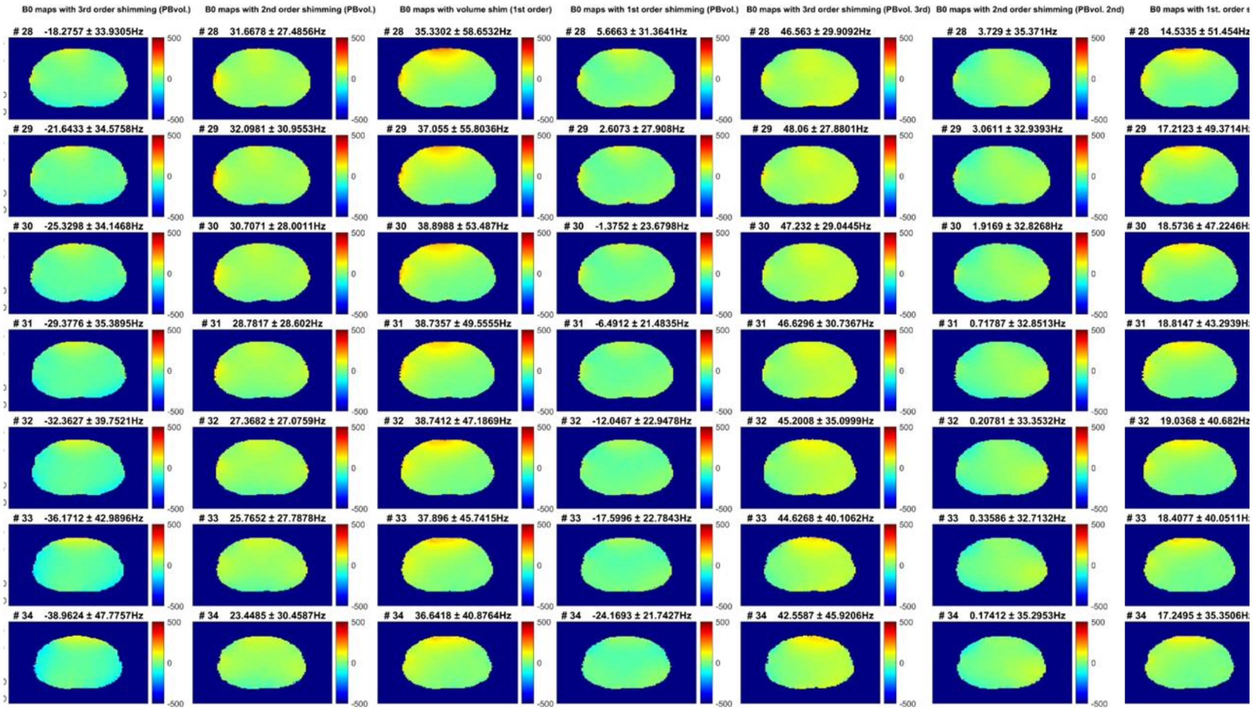
